# Genetic diversity analysis of the D614G mutation in SARS-CoV-2

**DOI:** 10.1101/2020.10.30.362954

**Authors:** Pierre Teodósio Felix, Dallynne Bárbara Ramos Venâncio, Eduarda Doralice Alves Braz Da Silva, Robson da Silva Ramos

**Author notes:** Corresponding author/Contact.

## Abstract

In this work, we evaluated the levels of genetic diversity in 18 genomes of SARS-CoV-2 carrying the D614G mutation, coming from Malaysia and Venezuela and publicly available at the National Center of Biotechnology and Information (NCBI). These haplotypes were previously used for phylogenetic analysis, following the LaBECom protocols. All gaps and unconserved sites were extracted for the construction of a phylogenetic tree. As specific methodologies for paired FST estimators, Molecular Variance (AMOVA), Genetic Distance, mismatch, demographic and spatial expansion analyses, molecular diversity and evolutionary divergence time analyses, 20,000 random permutations were always used. The results revealed the presence of only 57 sites of polymorphic and parsimonium-informative among the 29,827bp analyzed and the analyses based on F_ST_ values confirmed the presence of two distinct genetic entities with fixation index of 22% and with a higher component of population variation (78.14%). Tau variations revealed a significant time of divergence, supported by mismatch analysis of the observed distribution (τ = 42%). It is safe to say that the small number of existing polymorphisms should not reflect major changes in the protein products of viral populations in both countries and this consideration provides the safety that, although there are differences in the haplotypes studied, these differences are minimal for both regions analyzed geographically and, therefore, it seems safe to extrapolate the levels of polymorphism and molecular diversity found in the samples for other mutant genomes of SARS-CoV-2 in other countries. This reduces speculation about the possibility of large differences between mutant strains of SARS-CoV-2 (D614G) and wild strains, at least at the level of their protein products, although the mutant form has higher transmission speed and infection. The analyses suggest that possible variations in protein products, of the wild virus in relation to its mutant form, should be minimal, bringing peace of mind as to the increased risk of death from the new form of the virus, as well as possible problems of gradual adjustments in some molecular targets for vaccines.

## Introduction

The rapid transmission rate of SARS-CoV-2, detected in Wuhan (Hubei Province, China) in late 2019 resulted in the spread of the virus worldwide and in a few months. (MOHAMMAD *et al*, 2020) and even assuming that, despite its high transmissibility rate, the virus has a slow replication capability of its RNA, a variant with a single residual substitution called (D614G), it emerged as a dominant strain worldwide. (ZHANG *et al*, 2020).

The sequencing of SARS-CoV-2 showed that this mutation is already common in several continents and that it is always available, in only two super-clades (S and G), having its clinical parameters of severity or lethality, always associated with only one of them (S). (ELIZONDO *et al*, 2020). Phylogenetic analyses of some Sequences of SARS-CoV-2 from Uruguay revealed that there is a relative prevalence of variant S over G and at this time, the S-clate is already frequently found in Europe, reinforcing the indication that it is even more infectious than WT. (CASTILLO *et al*, 2020)

The D614G mutation, which is a change from a negatively charged aspartate (D) to glycine (G) in the loop region of the S1 domain of protein S (MOHAMMAD *et al*, 2020), as it progressed slowly in East Asia, would only appear at the end of April in viral samples of Chinese patients (90-97%), then described as arising at position 614 of protein S and resulting from a single change from nucleotide A to G at position 23,403 in the reference genome Wuhan-Hu-1, conferring structural advantages for the virus and an increase in its transmissibility and viral loads, its transduction in human cells and its pathogenicity and lethality, leading to speculation, including, that the efficacy of vaccines and countermeasures aimed at protein S, could be adversely affected, requiring a frequent correction of these vaccines or these countermeasures. (MCAULEY *et al*,2020)

Today, with a specific emergency pattern beyond China, the initial SARS-CoV-2 viral isolates are in different groups than those evolved later, such as SARS-CoV-2 isolates from India, which are clearly dispersed and that only the initial samples of Kerala are similar to wuhan isolates, all of which are predominantly originating from patients with a history of travel to Italy. This makes it clear that the mutation occurred mainly when the virus began to spread during the second acute infection and morbidity in Italy, Spain and other European countries. (GUPTA *et al*, 2020).

The exact impact of the mutation on the phenotype of the disease is a critical issue and still needs to be accurately elucidated. Combined genetic, structural and epidemiological data suggest that the D614G key may cause an increased prevalence of chemosensory deficits, as observed during the progression of the East Asian pandemic to Western countries. Increased binding of the G614 peak variant to the ACE2 host receptor and/or increased cell input efficiency by peak protein stabilization and reduced cleavage may be the underlying mechanism. (BUTOWT *et al*, 2020).

The global spread and increased infection of the SARS-CoV-2 variant, D614G raises the question of whether this structural change would compromise the efficacy of antiviral therapies directed to protein S, especially if they were designed to target D614. (YURKOVETSKIY *et al*, 2020).

Further studies are needed to determine how the growing diversity of mutation patterns influence the fitness and reproduction of viral populations, and how their susceptibility and avoidance to immune responses as well as treatments takes place. Thinking about it, the team from the Laboratory of Population Genetics and Computational Evolutionary Biology (LaBECom-UNIVISA) conducted a molecular variance study in 18 Mutant D614G haplotypes of SARS-CoV-2, from Malaysia and Venezuela, available at the National Center for Biotechnology Information (NCBI), to evaluate the possible levels of genetic diversity and polymorphisms existing in this particular PopSet.

## Objective

Evaluate possible levels of genetic diversity and polymorphisms existing in 18 mutant D614G haplotypes of SARS-CoV-2

## Methodology

1. **Databank**: 18 sequences of the complete genome of the SARS-CoV-2 virus carrying the D614G mutation and from Malaysia and Venezuela (16 and 2 haplotypes respectively), all with 29,827pb extension, were recovered from GENBANK (https://www.ncbi.nlm.nih.gov/nuccore/?term=D614G) on October 27, 2020. Once aligned using the MEGA X program (TAMURA *et al*., 2018), ambiguous sites, lost data and gaps, were excluded resulting in an analysisable sequence with only 57pb extension.
2. **For Viewing variable sites**: The graphical representation of the sites was made using the WEBLOGO v3 software, descrito por CROOKS *et al*., 2004.
3. **Genetic Structuring Analyses**: Paired F_ST_ estimators, Molecular Variance (AMOVA), Genetic Distance, mismatch, demographic and spatial expansion analyses, molecular diversity and evolutionary divergence time were obtained with the Software Arlequin v. 3.5 (EXCOFFIER *et al*., 2005) using 1000 random permutations (NEI and KUMAR, 2000). The FST and geographic distance matrices were not compared. All steps of this process are described below:

### Genetic diversity

Among the routines of LaBECom, this test is used to measure the genetic diversity that is equivalent to the heterozygosity expected in the groups studied. We used for this the standard index of genetic diversity H, described by Nei (1987). Which can also be estimated by the method proposed by PONS and PETIT (1995).

#### Site frequency spectrum (SFS)

According to LaBECom protocols, we used this local frequency spectrum analytical test (SFS), from DNA sequence data that allows us to estimate the demographic parameters of the frequency spectrum. Simulations are made using fastsimcoal2 software, available in http://cmpg.unibe.ch/software/fastsimcoal2/.

#### Molecular diversity indices

Molecular diversity indices are obtained by means of the average number of paired differences, as described by Tajima in 1993, in this test we used sequences that do not fit the model of neutral theory that establishes the existence of a balance between mutation and genetic drift.

#### Calculating THETA estimators

Theta population parameters are used in our Laboratory when we want to qualify the genetic diversity of the populations studied. These estimates, classified as Theta Hom – which aim to estimate the expected homozygosity in a population in equilibrium between drift and mutation and the estimates Theta (S) (WATTERSON, 1975), Theta (K) (EWENS, 1972) and Theta (π) (TAJIMA, 1983).

#### Calculation of the distribution of mismatch

In LaBECom, analyses of the mismatch distribution are always performed relating the observed number of differences between haplotype pairs, trying to define or establish a pattern of population demographic behavior, as already described by (ROGERS; HARPENDING, 1992; Hudson, Hudson, HUDSON, SLATKIN, 1991; RAY et al., 2003, EXCOFFIER, 2004).

#### Pure demographic expansion

This model is always used when we intend to estimate the probability of differences observed between two haplotypes not recombined and randomly chosen, this methodology in our laboratory is used when we assume that the expansion, in a haploid population, reached a momentary balance even having passed through τ generations, of sizes 0 N to 1 N. In this case, the probability of observing the S differences between two non-recombined and randomly chosen haplotypes is given by the probability of observing two haplotypes with S differences in this population (Watterson, 1975).

#### Spatial expansion

The use of this model in LaBECom is usually indicated if the reach of a population is initially restricted to a very small area, and when one notices signs of a growth of the same, in the same space and over a relatively short time. The resulting population generally becomes subdivided in the sense that individuals tend to mate with geographically close individuals rather than random individuals. To follow the dimensions of spatial expansion, we at LaBECom always take into account:

L: Number of loci
Gamma Correction: This fix is always used when mutation rates do not seem uniform for all sites.
nd: Number of substitutions observed between two DNA sequences.
ns: Number of transitions observed between two DNA sequences.
nv: Number of transversions observed between two DNA sequences.
ω: G + C ratio, calculated in all DNA sequences of a given sample.
Paired Difference: Shows the number of loci for which two haplotypes are different.
Percentage difference: This difference is responsible for producing the percentage of loci for which two haplotypes are different.

#### Haplotypic inferences

We use these inferences for haplotypic or genotypic data with unknown gametic phase. Following our protocol, inferences are estimated by observing the relationship between haplotype i and xi times its number of copies, generating an estimated frequency (^pi). With genotypic data with unknown gametic phase, the frequencies of haplotypes are estimated by the maximum likelihood method, and can also be estimated using the expected Maximization (MS) algorithm.

#### Method of Jukes and Cantor

This method, when used in LaBECom, allows estimating a corrected percentage of how different two haplotypes are. This correction allows us to assume that there have been several substitutions per site, since the most recent ancestor of the two haplotypes studied. Here, we also assume a correction for identical replacement rates for all four nucleotides A C, G and T.

#### Kimura method with two parameters

Much like the previous test, this fix allows for multiple site substitutions, but takes into account different replacement rates between transitions and transversions.

#### Tamura method

We at LaBECom understand this method as an extension of the 2-parameter Kimura method, which also allows the estimation of frequencies for different haplotypes. However, transition-transversion relationships as well as general nucleotide frequencies are calculated from the original data.

#### Tajima and Nei method

At this stage, we were also able to produce a corrected percentage of nucleotides for which two haplotypes are different, but this correction is an extension of the Jukes and Cantor method, with the difference of being able to do this from the original data.

#### Tamura and Nei model

As in kimura’s models 2 parameters a distance of Tajima and Nei, this correction allows, inferring different rates of transversions and transitions, besides being able to distinguish transition rates between purines and pyrimidines.

#### Minimum spanning network

To calculate the distance between OTU (operational taxonomic units) from the paired distance matrix of haplotypes, we used a Minimum Spanning Network (MSN) tree, with a slight modification of the algorithm described in Rohlf (1973). We usually use free software written in Pascal called MINSPNET. EXE running in DOS language, previously available at: http://anthropologie.unige.ch/LGB/software/win/min-span-net/.

#### Genotypic data with unknown gametic phase

##### EM algorithm

To estimate haplotypic frequencies we used the maximum likelihood model with an algorithm that maximizes the expected values. The use of this algorithm in LaBECom, allows to obtain the maximum likelihood estimates from multilocal data of gamtic phase is unknown (phenotypic data). It is a slightly more complex procedure since it does not allow us to do a simple gene count, since individuals in a population can be heterozygous to more than one locus.

##### ELB algorithm

Very similar to the previous algorithm, ELB attempts to reconstruct the gametic phase (unknown) of multilocal genotypes by adjusting the sizes and locations of neighboring loci to explore some rare recombination.

#### Neutrality tests

##### Ewens-Watterson homozygosis test

We use this test in LaBECom for both haploid and diploid data. This test is used only as a way to summarize the distribution of allelic frequency, without taking into account its biological significance. This test is based on the sampling theory of neutral alllinks from Ewens (1972) and tested by Watterson (1978). It is now limited to sample sizes of 2,000 genes or less and 1,000 different alleles (haplotypes) or less. It is still used to test the hypothesis of selective neutrality and population balance against natural selection or the presence of some advantageous alleles.

##### Accurate Ewens-Watterson-Slatkin Test

This test created by Slatikin in 1994 and adapted by himself in 1996. is used in our protocols when we want to compare the probabilities of random samples with those of observed samples.

##### Chakraborty’s test of population amalgamation

This test was proposed by Chakrabordy in 1990, serves to calculate the observed probability of a randomly neutral sample with a number of alleles equal to or greater than that observed, it is based on the infinite allele model and sampling theory for neutral Alleles of Ewens (1972).

##### Tajima Selective Neutrality Test

We use this test in our Laboratory when DNA sequences or haplotypes produced by RFLP are short. It is based on the 1989 Tajima test, using the model of infinite sites without recombination. It commutes two estimators using the theta mutation as a parameter.

##### *FS FU* Test of Selective Neutrality

Also based on the model of infinite sites without recombination, the FU test is suitable for short DNA sequences or haplotypes produced by RFLP. However, in this case, it assesses the observed probability of a randomly neutral sample with a number of alleles equal to or less than the observed value. In this case the estimator used is θ.

#### Methods that measure interpopulation diversity

##### Genetic structure of the population inferred by molecular variance analysis (AMOVA)

This stage is the most used in the LaBECom protocols because it allows to know the genetic structure of populations measuring their variances, this methodology, first defined by Cockerham in 1969 and 1973) and, later adapted by other researchers, is essentially similar to other approaches based on analyses of gene frequency variance, but takes into account the number of mutations between haplotypes. When the population group is defined, we can define a particular genetic structure that will be tested, that is, we can create a hierarchical analysis of variance by dividing the total variance into covariance components by being able to measure intra-individual differences, interindividual differences and/or interpopulation allocated differences.

##### Minimum Spanning Network (MSN) among haplotypes

In LaBECom, this tree is generated using the operational taxonomic units (OTU). This tree is calculated from the matrix of paired distances using a modification of the algorithm described in Rohlf (1973).

##### Locus-by-locus AMOVA

We performed this analysis for each locus separately as it is performed at the haplotypic level and the variance components and f statistics are estimated for each locus separately generating in a more global panorama.

##### Paired genetic distances between populations

This is the most present analysis in the work of LaBECom. These generate paired FST parameters that are always used, extremely reliably, to estimate the short-term genetic distances between the populations studied, in this model a slight algorithmic adaptation is applied to linearize the genetic distance with the time of population divergence (Reynolds et al. 1983; Slatkin, 1995).

##### Reynolds Distance (Reynolds et al. 1983)

Here we measured how much pairs of fixed N-size haplotypes diverged over t generations, based on F_ST_ indices.

##### Slatkin’s linearized F_ST’s_ (Slatkin 1995)

We used this test in LaBECom when we want to know how much two Haploid populations of N size diverged t generations behind a population of identical size and managed to remain isolated and without migration. This is a demographic model and applies very well to the phylogeography work of our Laboratory.

##### Nei’s average number of differences between populations

In this test we assumed that the relationship between the gross (D) and liquid (AD) number of Nei differences between populations is the increase in genetic distance between populations (Nei and Li, 1979).

##### Relative population sizes: divergence between populations of unequal sizes

We used this method in LaBECom when we want to estimate the time of divergence between populations of equal sizes (Gaggiotti and Excoffier, 2000), assuming that two populations diverged from an ancestral population of N0 size a few t generations in the past, and that they have remained isolated from each other ever since. In this method we assume that even though the sizes of the two child populations are different, the sum of them will always correspond to the size of the ancestral population. The procedure is based on the comparison of intra and inter populational (π’s) diversities that have a large variance, which means that for short divergence times, the average diversity found within the population may be higher than that observed among populations. These calculations should therefore be made if the assumptions of a pure fission model are met and if the divergence time is relatively old. The results of this simulation show that this procedure leads to better results than other methods that do not take into account unequal population sizes, especially when the relative sizes of the daughter populations are in fact unequal.

##### Accurate population differentiation tests

We at LaBECom understand that this test is an analog of fisher’s exact test in a 2×2 contingency table extended to a rxk contingency table. It has been described in Raymond and Rousset (1995) and tests the hypothesis of a random distribution of k different haplotypes or genotypes among r populations.

##### Assignment of individual genotypes to populations

Inspired by what had been described in Paetkau et al (1995, 1997) and Waser and Strobeck (1998) this method determines the origin of specific individuals, knowing a list of potential source populations and uses the allelic frequencies estimated in each sample from their original constitution.

##### Detection of loci under selection from F-statistics

We use this test when we suspect that natural selection affects genetic diversity among populations. This method was adapted by Cavalli-Sforza in 1996 from a 1973 work by Lewontin and Krakauer.

#### Results

All 18 sequences of the Mutant SARS-CoV-2 (D614G) virus from Malaysia and Venezuela (Table 1), revealed the presence of only 57 polymorphic and informative parsimony sites among the 29,827bp analyzed. The graphical representation of these sites can be seen in a logo built with the PROGRAM WEBLOGO v 3.7.4., where the size of each nucleotide is proportional to its frequency for the certain sites. (Figure 1).

**Table 1.**
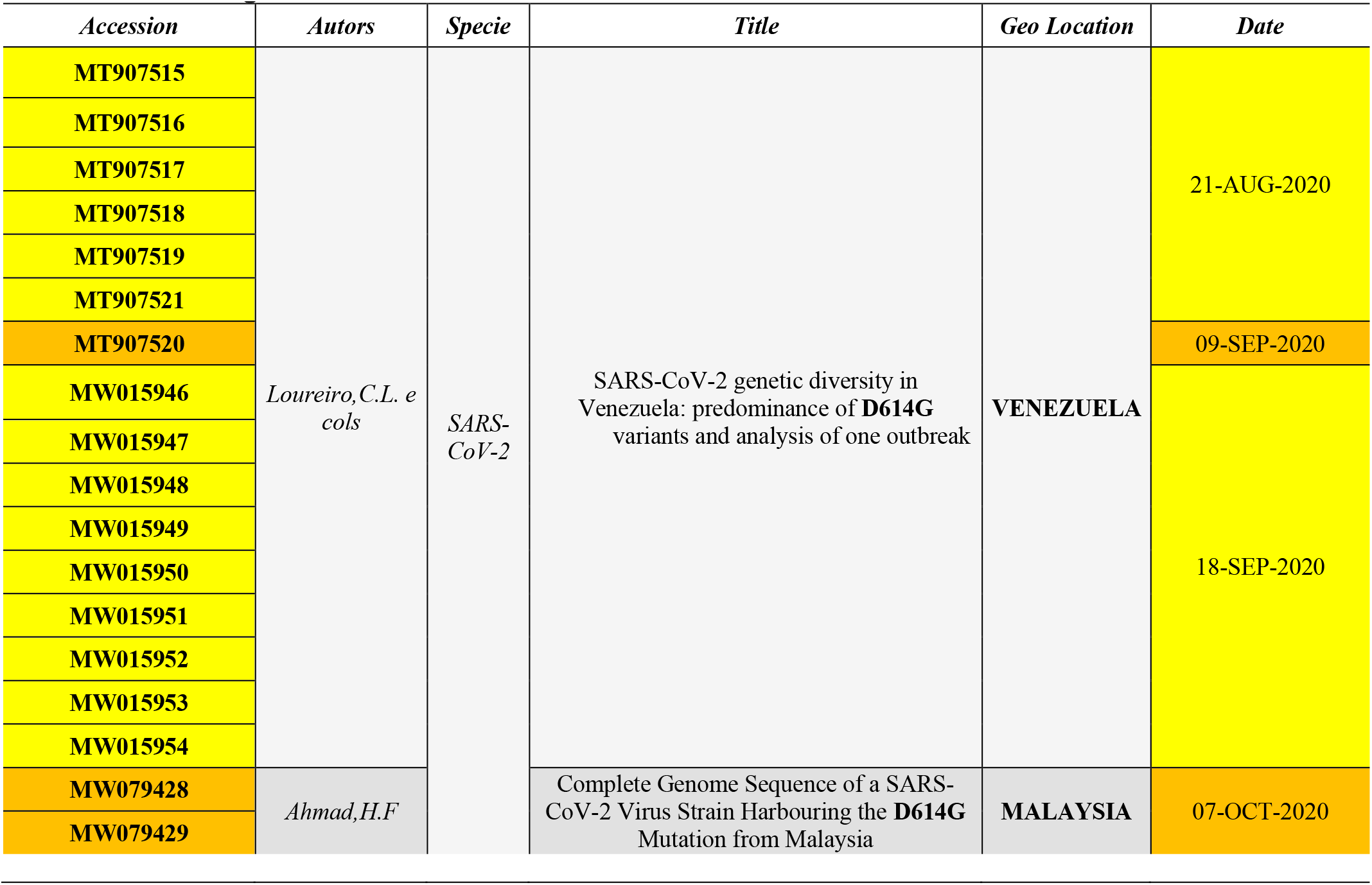
Description of all sequences analyzed in this work. By GENBANK access number, authors, Geo Location and filing date.

**Figure 1.**
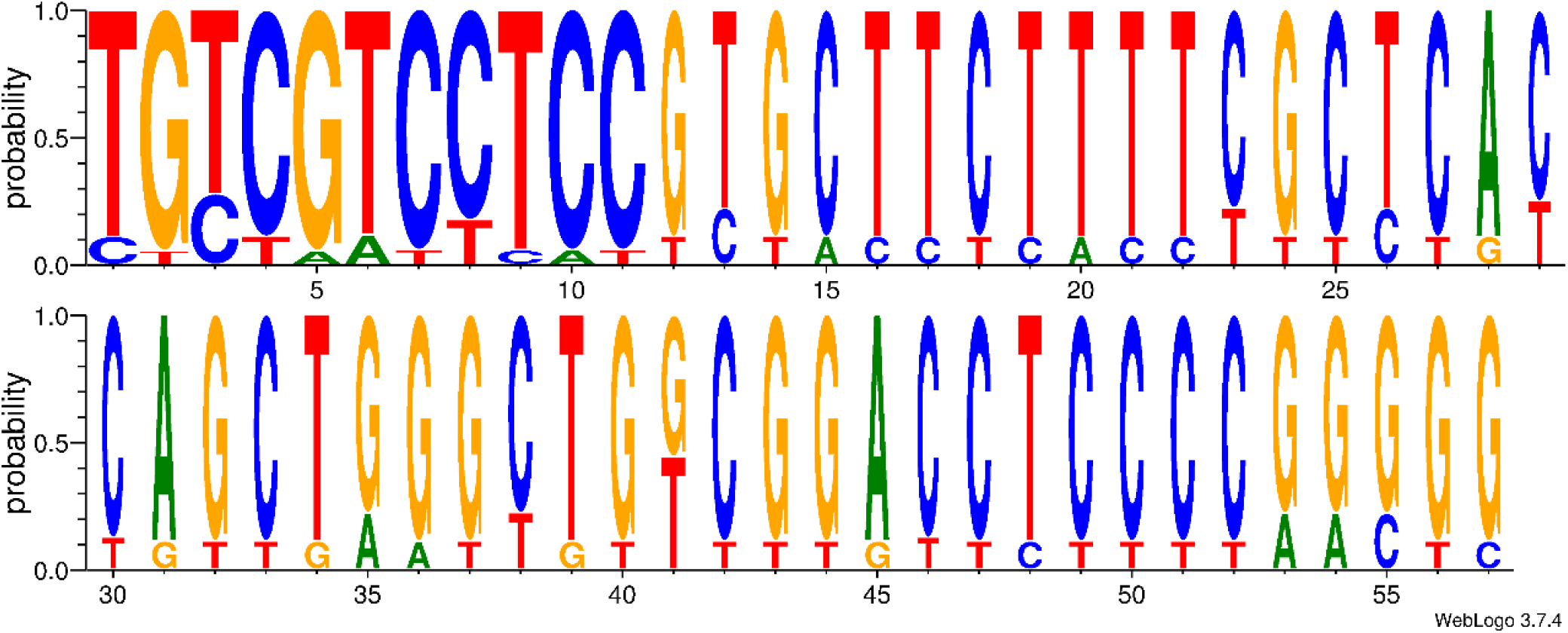
Graphic representation of the 57 parsimonium-informative sites of SARS-CoV-2 carriers of the D614G mutation.

Using the UPGMA method, for the 57 parsimony-informative sites, it was possible to understand that the 18 haplotypes comprised two distinct groups, and it is even possible that there are haplotype sharing between the countries studied (Figure 2).

**Figure 2.**
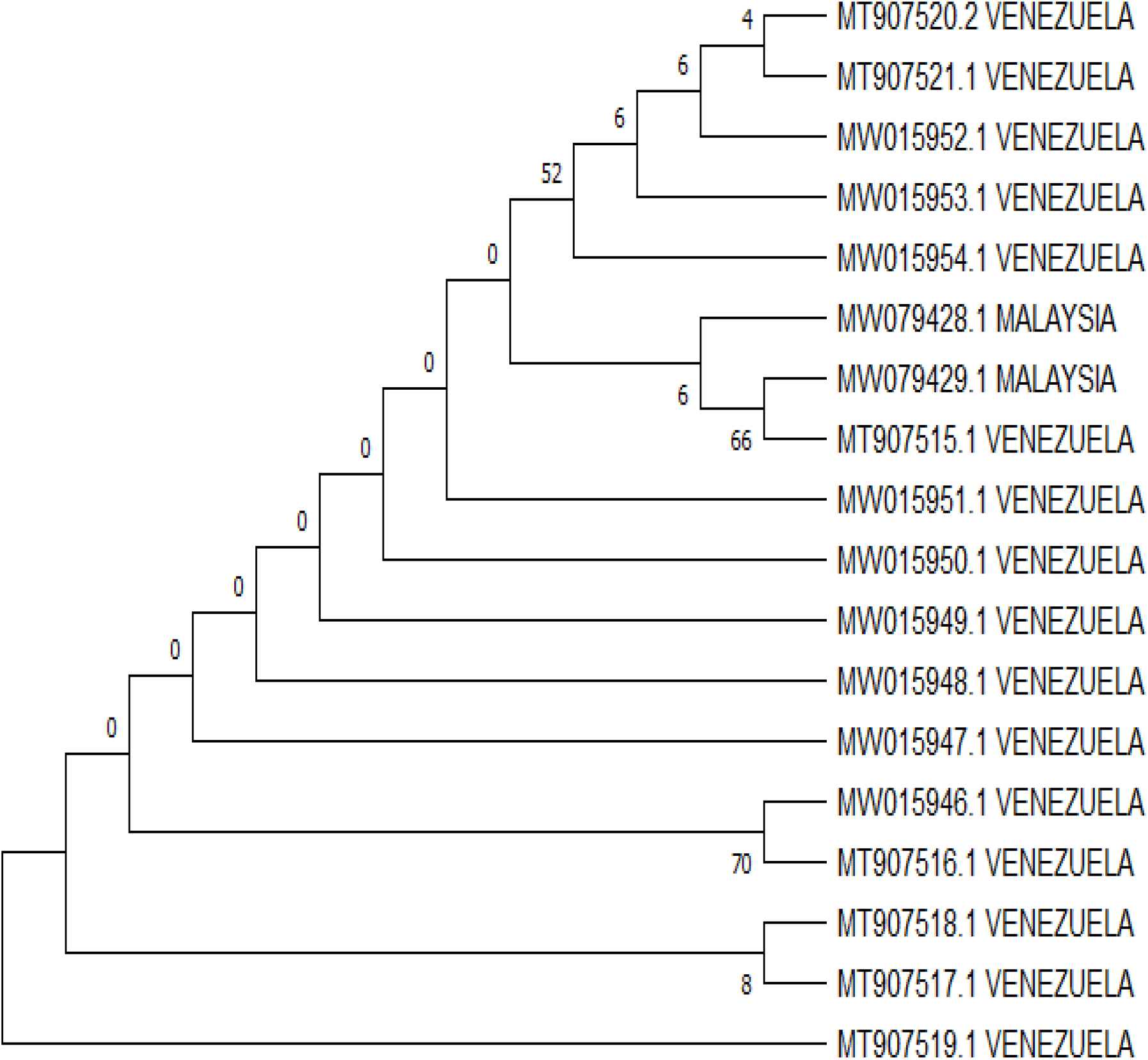
Evolutionary analysis by the maximum likelihood method. The evolutionary history was inferred using the maximum likelihood method and the 3-parameter Tamura model [1]. The tree with the highest probability of logging (−1366.35) is shown. The percentage of trees in which the associated dollar sums group together is shown next to the branches. The initial trees for heuristic research were obtained automatically by applying the Join-Join and BioNJ algorithms to an array of distances in estimated pairs using the Tamura 3 parameter model and then selecting the topology with a higher log probability value. This analysis involved 38 nucleotide sequences. The evolutionary analyses were performed in MEGA X.

## Molecular Variance Analysis (AMOVA) and Genetic Distance

Analyses based on FST values confirmed the presence of two distinct genetic entities with a fixation index of 22% and with a larger component of within populations variation (78.14%)***p*** less than 0.05. Significant evolutionary divergences were observed within and between the groups (Table 1) and a high genetic similarity between the sequences that made up the Malaysia group, as well as a greater evolutionary divergence between the sequences that made up the Group of Venezuela (Table 2, Table 3, Figure 3 and Figure 4).

**Table 2.** Components of haplotypic variation and paired F_ST_ value of the parsimonium-informative sites of SARS-CoV-2 carriers of the D614G mutation.

| SOURCE OF VARIATION | D.F. | SUM OF SQUARES | VARIANCE COMPONENTS | PERCENTAGE OF VARIATION |
| --- | --- | --- | --- | --- |
| Among populations | 1 | 26.410 | 3.70447 Va | 21.86 |
| Within populations | 16 | 211.812 | 13.23828 Vb | 78.14 |
| Total | 17 | 238.222 | 16.94275 |  |
| **Fixation Index $F_{ST}$: 0.21865**<br>Significance tests (110 permutations)<br>Va and $F_{ST}$: P (rand. value > obs. value) = 0.19091<br>P (rand. value = obs. value) = 0.02727<br>P-value = 0.21818+-0.05107 | | | | |

**Table 3.**
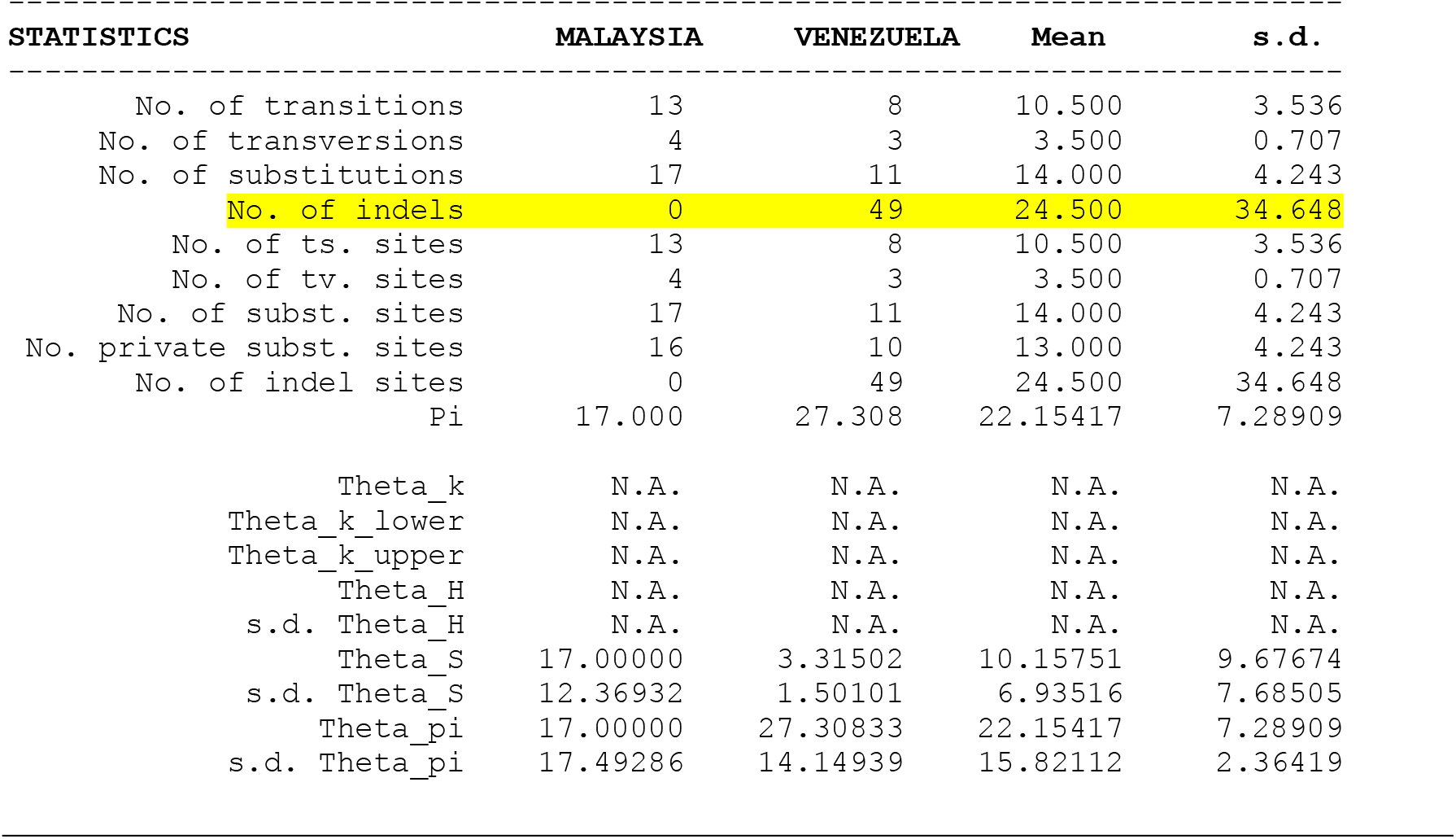
Molecular Diversity Indices of the parsimonium-informative sites of SARS-CoV-2 carriers of the D614G mutation.

| STATISTICS | MALAYSIA | VENEZUELA | Mean | s.d. |
| --- | --- | --- | --- | --- |
| No. of transitions | 13 | 8 | 10.500 | 3.536 |
| No. of transversions | 4 | 3 | 3.500 | 0.707 |
| No. of substitutions | 17 | 11 | 14.000 | 4.243 |
| No. of indels | 0 | 49 | 24.500 | 34.648 |
| No. of ts. sites | 13 | 8 | 10.500 | 3.536 |
| No. of tv. sites | 4 | 3 | 3.500 | 0.707 |
| No. of subst. sites | 17 | 11 | 14.000 | 4.243 |
| No. private subst. sites | 16 | 10 | 13.000 | 4.243 |
| No. of indel sites | 0 | 49 | 24.500 | 34.648 |
| Pi | 17.0000 | 27.308 | 22.15417 | 7.28909 |
| Theta_k | N.A. | N.A. | N.A. | N.A. |
| Theta_k_lower | N.A. | N.A. | N.A. | N.A. |
| Theta_k_upper | N.A. | N.A. | N.A. | N.A. |
| Theta_H | N.A. | N.A. | N.A. | N.A. |
| s.d. Theta_H | N.A. | N.A. | N.A. | N.A. |
| Theta_S | 17.00000 | 3.31502 | 10.15751 | 9.67674 |
| s.d. Theta_S | 12.36932 | 1.50101 | 6.93516 | 7.68505 |
| Theta_pi | 17.00000 | 27.30833 | 22.15417 | 7.28909 |
| s.d. Theta_pi | 17.49286 | 14.14939 | 15.82112 | 2.36419 |

**Figure 3.**
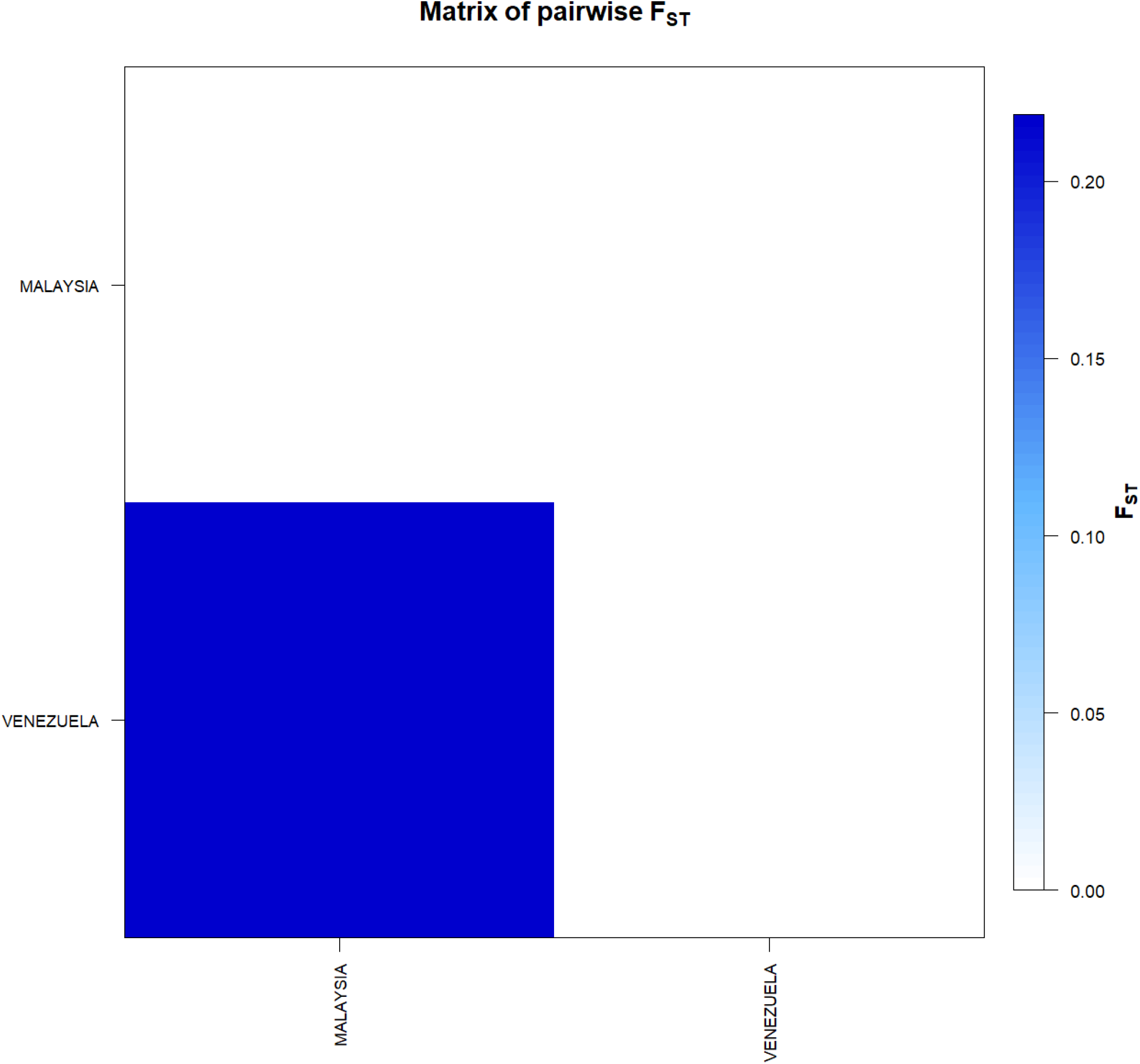
F_ST_-based genetic distance matrix between for the sequences of SARS-CoV-2 carriers of the D614G mutation. * Generated by the statistical package in R language using the output data of the Software Arlequin version 3.5.1.2

**Figure 4.**
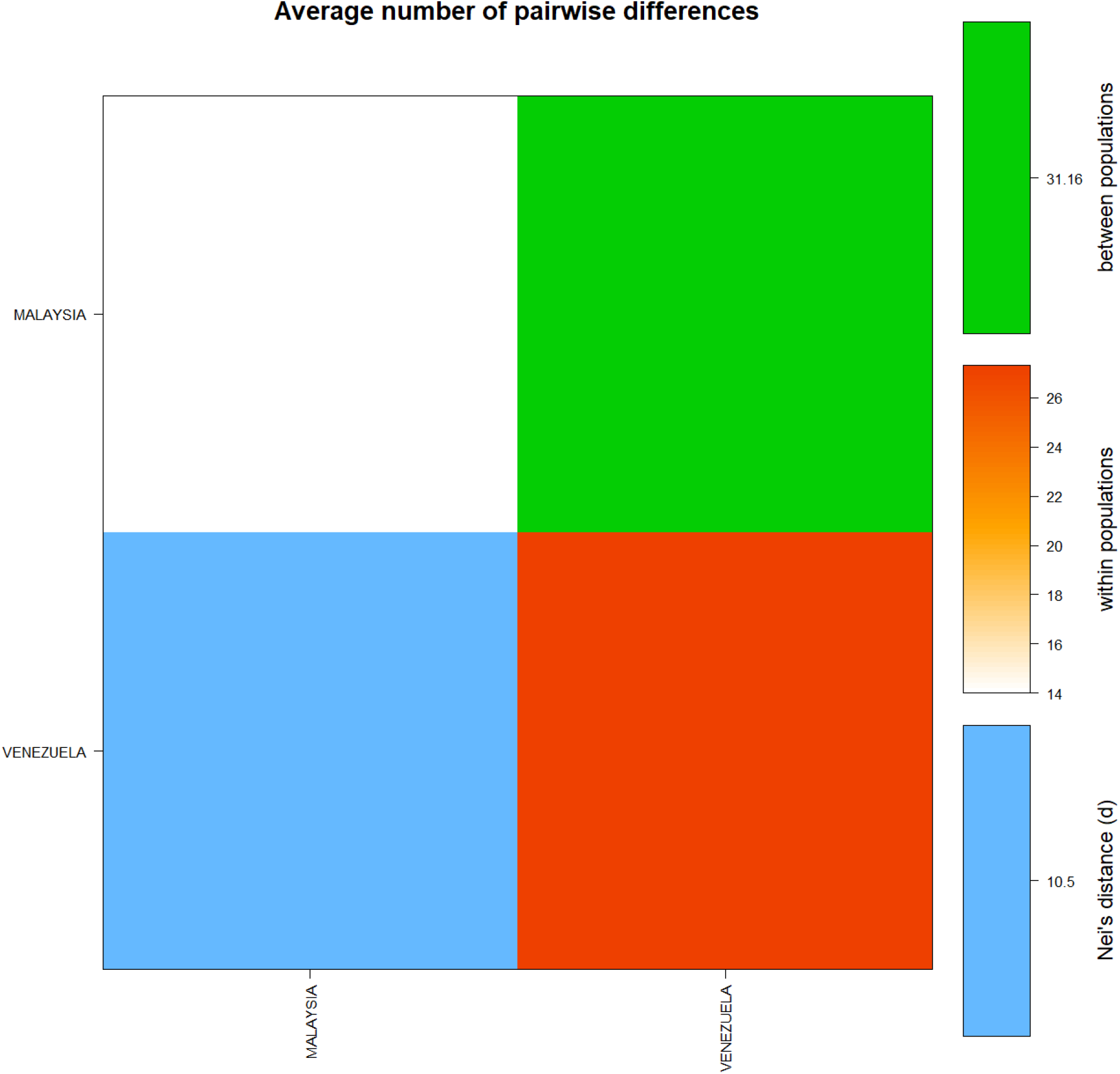
Matrix of paired differences between the populations studied: between the groups; within the groups; and Nei distance for the sequences of SARS-CoV-2 carriers of the D614G mutation.

The *Tau* variations (related to the ancestry of the two groups) revealed a significant time of divergence, supported by the mismatch analysis of the observed distribution (τ = 42%) and with constant mutation rates between localities (Table 4).

**Table 4.** Tau Values (τ) for the sequences of SARS-CoV-2 carriers of the D614G mutation.

| POPULATIONS | MALAYSIA | VENEZUELA |
| --- | --- | --- |
| MALAYSIA | 0.00000 | **3.34198** |
| VENEZUELA | **3.34198** | 0.00000 |

**Figure 5.**
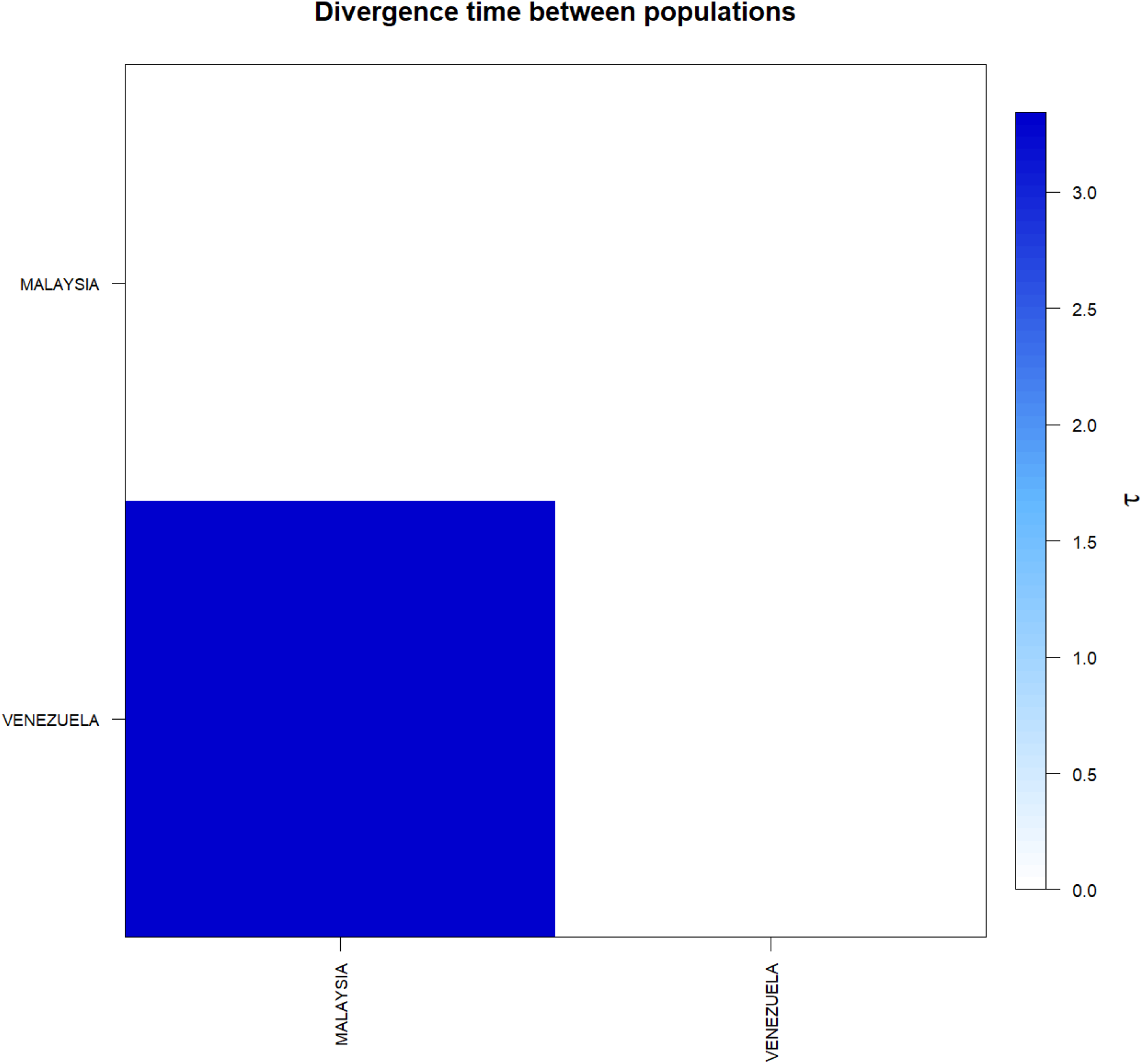
Matrix of divergence time between for the sequences of SARS-CoV-2 carriers of the D614G mutation. In evidence the high value τ present between the sequences of Brazil and Venezuela. * Generated by the statistical package in R language using the output data of the Software Arlequin version 3.5.1.2.

The molecular analyses estimated by θ reflected a significant level of mutations among all haplotypes (transitions and transversions). Indels mutations (insertions or additions), were found only in the group of haplotypes from Venezuela (Table 3) (Figure 8). The Tajima and Fs de Fu tests showed disagreements between the estimates of general φ and π, but with negative and highly significant values, indicating, once again, the absence of population expansions in the Malaysia group (Table 5). The irregularity index (R= Raggedness) with parametric bootstrap simulated new values φ for before and after a supposed demographic expansion and, in this case, assumed a value equal to zero for only the Malaysia group, probably due to the small number of haplotypes analyzed (Table 6); (Figure 6 and figure 7).

**Table 5.** Neutrality Tests for the sequences of SARS-CoV-2 carriers of the D614G mutation.

|  | Statistics | MALAYSIA | VENEZUELA | Mean | s.d. |
| --- | --- | --- | --- | --- | --- |
| **Ewens-Watterson test** |  |  |  |  |  |
| ----- |  |  |  |  |  |
|  | Sample size | 2 | 16 | 9.00000 | 9.89949 |
|  | No. of alleles (unchecked) | 2 | 16 | 9.00000 | 9.89949 |
|  | Observed F value | N.A. | N.A. | N.A. | N.A. |
|  | Expected F value | N.A. | N.A. | N.A. | N.A. |
| | Watterson test: $\Pr(\text{rand } F \leq \text{obs } F)$ | N.A. | N.A. | N.A. | N.A. |
|  | Slatkin's exact test P-value | N.A. | N.A. | N.A. | N.A. |
| **Chakraborty's test** |  |  |  |  |  |
| ----- |  |  |  |  |  |
|  | Sample size | 2 | 16 | 9.00000 | 9.89949 |
|  | No. of alleles (unchecked) | 2 | 16 | 9.00000 | 9.89949 |
|  | Obs. homozygosity | 0.00000 | 0.00000 | 0.00000 | 0.00000 |
|  | Exp. no. of alleles | 1.94444 | 12.77988 | 7.36216 | 7.66181 |
|  | P(k or more alleles) | N.A. | N.A. | N.A. | N.A. |
| **Tajima's D test** |  |  |  |  |  |
| ----- |  |  |  |  |  |
|  | Sample size | 2 | 16 | 9.00000 | 9.89949 |
|  | S | 17 | 11 | 14.00000 | 4.24264 |
|  | Pi | 17.00000 | 27.30833 | 22.15417 | 7.28909 |
|  | Tajima's D | 0.00000 | 0.76166 | 0.38083 | 0.53858 |
|  | Tajima's D p-value | 1.00000 | 0.78000 | 0.89000 | 0.15556 |
| **Fu's FS test** |  |  |  |  |  |
| ----- |  |  |  |  |  |
|  | No. of alleles (unchecked) | 2 | 16 | 9.00000 | 9.89949 |
|  | Theta_pi | 17.00000 | 27.30833 | 22.15417 | 7.28909 |
|  | Exp. no. of alleles | 1.94444 | 12.77988 | 7.36216 | 7.66181 |
|  | FS | 2.83321 | -3.71603 | -0.44141 | 4.63102 |
|  | FS p-value | 0.61000 | 0.07000 | 0.34000 | 0.38184 |

**Table 6.** Demographic and spatial expansion simulations based on the τ, θ, and M indices for the sequences of SARS-CoV-2 carriers of the D614G mutation.

| Statistics | MALAYSIA | VENEZUELA | Mean | s.d. |
| --- | --- | --- | --- | --- |
| **Demographic expansion** |  |  |  |  |
| Tau | 0.00000 | 0.00000 | 0.00000 | 0.00000 |
| Tau qt 2.5% | 0.00000 | 0.85937 | 0.42969 | 0.60767 |
| Tau qt 5% | 0.00000 | 1.06251 | 0.53126 | 0.75131 |
| Tau qt 95% | 0.00000 | 33.94924 | 16.97462 | 24.00573 |
| Tau qt 97.5% | 0.00000 | 38.66216 | 19.33108 | 27.33827 |
| Theta0 | 0.00000 | 57.59971 | 28.79985 | 40.72914 |
| Theta0 qt 2.5% | 0.00000 | 0.00000 | 0.00000 | 0.00000 |
| Theta0 qt 5% | 0.00000 | 0.00000 | 0.00000 | 0.00000 |
| Theta0 qt 95% | 0.00000 | 76.38712 | 38.19356 | 54.01385 |
| Theta0 qt 97.5% | 0.00000 | 83.87887 | 41.93943 | 59.31132 |
| Theta1 | 0.00000 | 3590.39020 | 1795.19510 | 2538.78926 |
| Theta1 qt 2.5% | 0.00000 | 37.46331 | 18.73166 | 26.49056 |
| Theta1 qt 5% | 0.00000 | 44.72598 | 22.36299 | 31.62604 |
| Theta1 qt 95% | 0.00000 | 520.91039 | 260.45519 | 368.33927 |
| Theta1 qt 97.5% | 0.00000 | 674.30700 | 337.15350 | 476.80706 |
| SSD | 0.00000 | 0.05955 | 0.02978 | 0.04211 |
| Model (SSD) p-value | 0.00000 | 0.00000 | 0.00000 | 0.00000 |
| Raggedness index | 0.00000 | 0.02951 | 0.01476 | 0.02087 |
| Raggedness p-value | 0.00000 | 0.35000 | 0.17500 | 0.24749 |
| **Spatial expansion** |  |  |  |  |
| Tau | 0.00000 | 41.09920 | 20.54960 | 29.06152 |
| Tau qt 2.5% | 0.00000 | 2.16582 | 1.08291 | 1.53147 |
| Tau qt 5% | 0.00000 | 3.91431 | 1.95716 | 2.76784 |
| Tau qt 95% | 0.00000 | 54.69081 | 27.34541 | 38.67225 |
| Tau qt 97.5% | 0.00000 | 59.33696 | 29.66848 | 41.95757 |
| Theta | 0.00000 | 12.18978 | 6.09489 | 8.61948 |
| Theta qt 2.5% | 0.00000 | 0.00072 | 0.00036 | 0.00051 |
| Theta qt 5% | 0.00000 | 0.08051 | 0.04025 | 0.05693 |
| Theta qt 95% | 0.00000 | 21.46667 | 10.73334 | 15.17923 |
| Theta qt 97.5% | 0.00000 | 23.06428 | 11.53214 | 16.30891 |
| M | 0.00000 | 2.15421 | 1.07711 | 1.52326 |
| M qt 2.5% | 0.00000 | 0.33692 | 0.16846 | 0.23824 |
| M qt 5% | 0.00000 | 0.58412 | 0.29206 | 0.41303 |
| M qt 95% | 0.00000 | 25.38039 | 12.69019 | 17.94664 |
| M qt 97.5% | 0.00000 | 38.24056 | 19.12028 | 27.04016 |
| SSD | 0.00000 | 0.04546 | 0.02273 | 0.03214 |
| Model (SSD) p-value | 0.00000 | 0.09000 | 0.04500 | 0.06364 |
| Raggedness index | 0.00000 | 0.02951 | 0.01476 | 0.02087 |
| Raggedness p-value | 0.00000 | 0.84000 | 0.42000 | 0.59397 |

**Figure 6.**
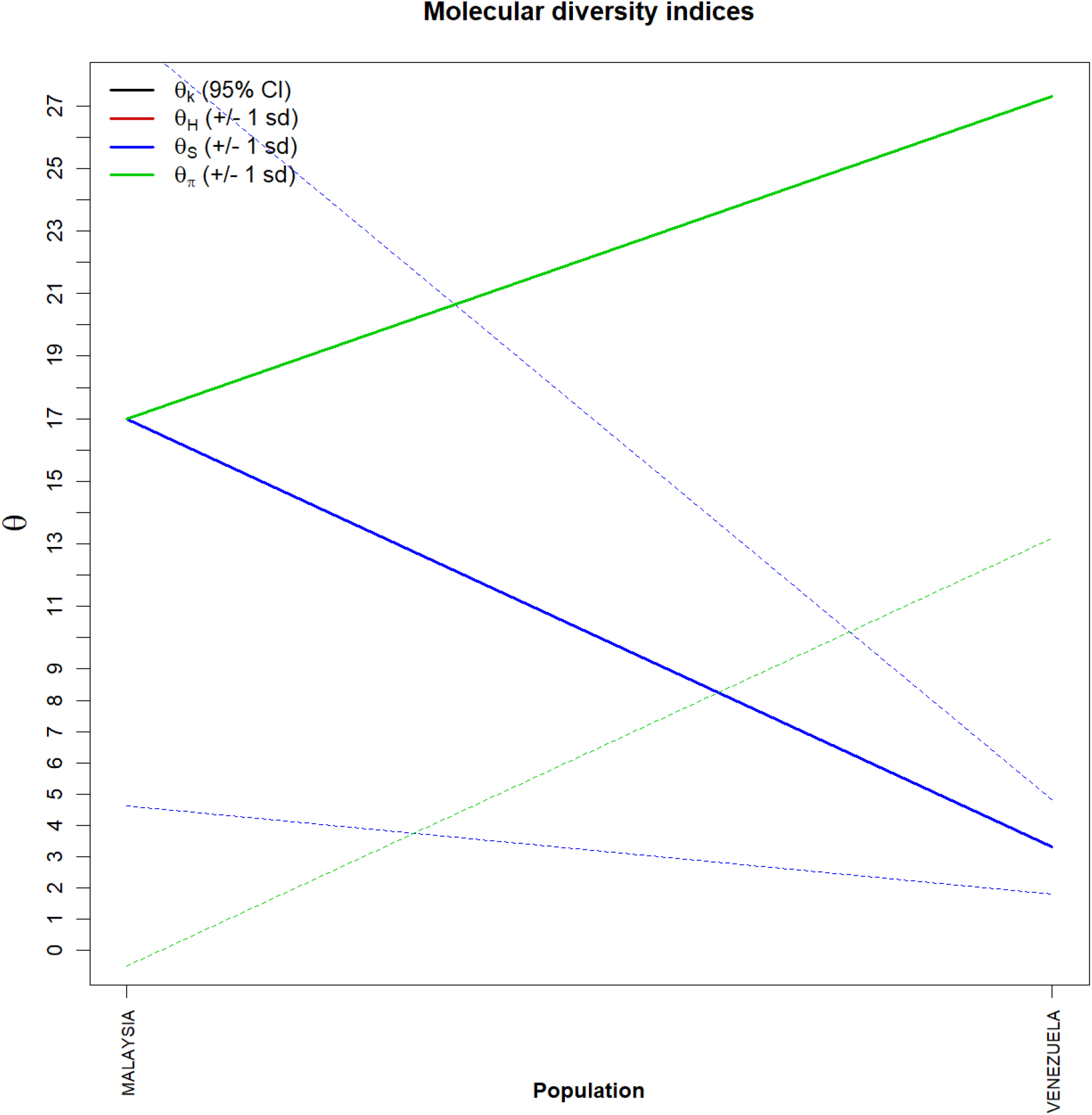
Graph of molecular diversity indices for the for the sequences of SARS-CoV-2 carriers of the D614G mutation. In the graph the values of θ: (θk) Relationship between the expected number of alllos (k) and the sample size; (θH) Expected homozygosity in a balanced relationship between drift and mutation; (θS) Relationship between the number of segregating sites (S), sample size (n) and non-recombinant sites; (θπ) Relationship between the average number of paired differences (π) and θ. * Generated by the statistical package in R language using the output data of the Arlequin software version 3.5.1.2.

**Figure 7.**
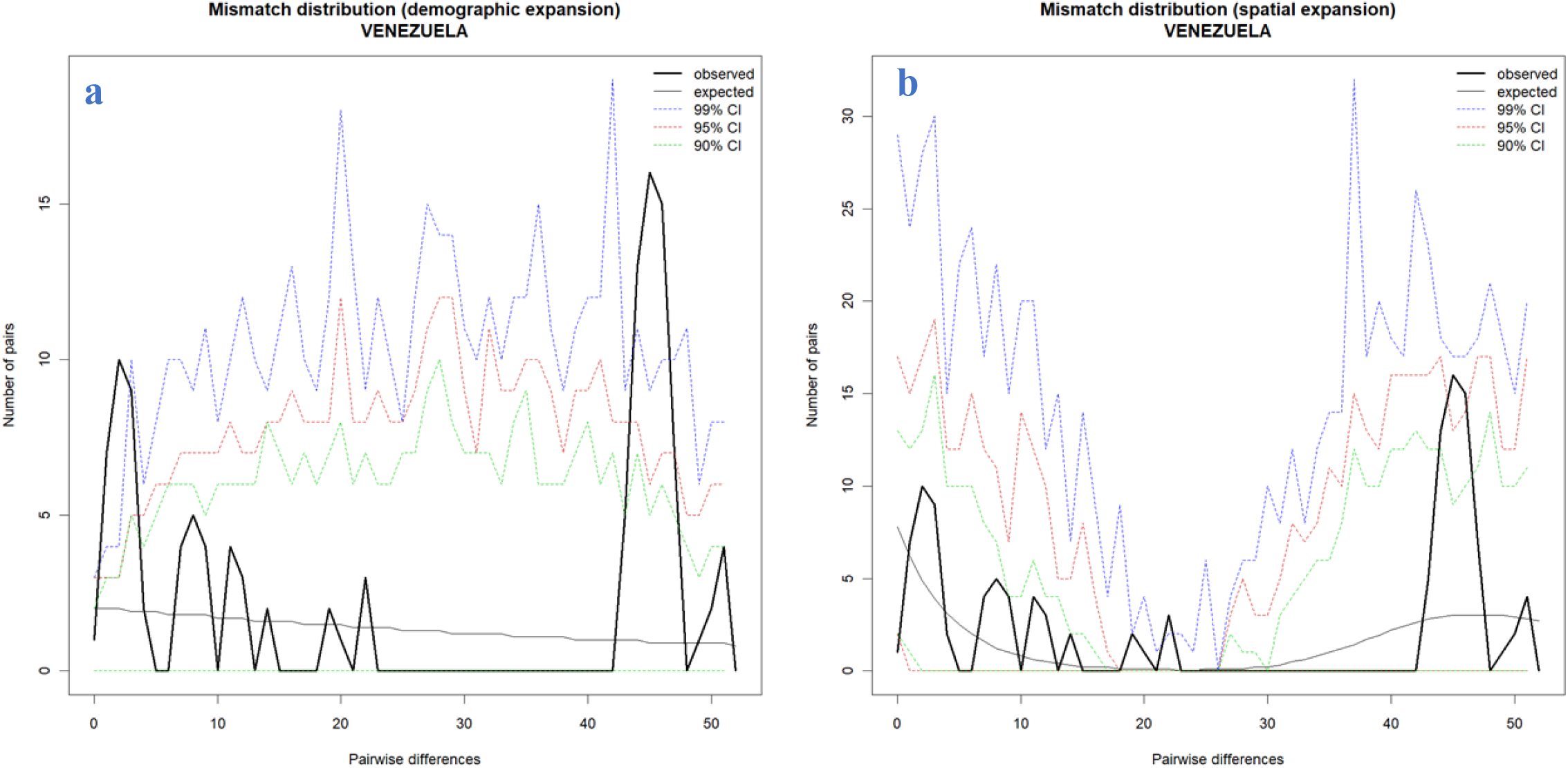
Comparison between the Demographic and Spatial Expansion for the for the sequences of SARS-CoV-2 carriers of the D614G mutation. (**a** and **b**) Graphs of demographic expansion and spatial expansion of haplotypes from Venezuela. *Graphs Generated by the statistical package in R language using the output data of the Software Arlequin version 3.5.1.2

**Figure 8.**
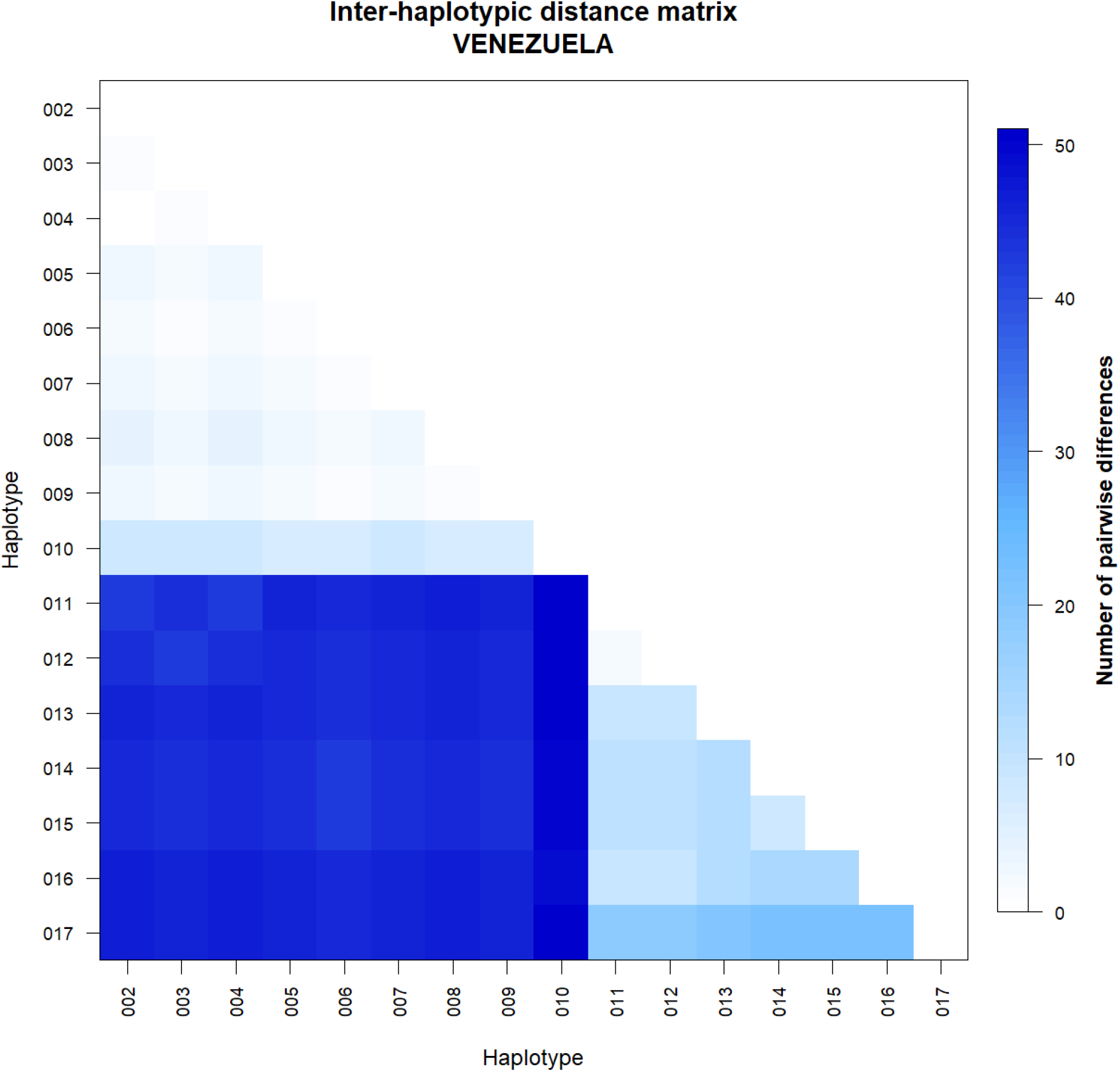
Matrix of inter haplotypic distance in the sequences of SARS-CoV-2 carriers of the D614G mutation from Venezuela. **Note the great variation between haplotypes**. *Generated by the statistical package in R language using the output data of the Software Arlequin version 3.5.1.2.

A significant similarity was also evidenced for the time of genetic evolutionary divergence among all populations; supported by τ variations, mismatch analyses and demographic and spatial expansion analyses. With a representative exception for haplotypes from Venezuela (Figures 8 and figure 9).

**Figure 9.**
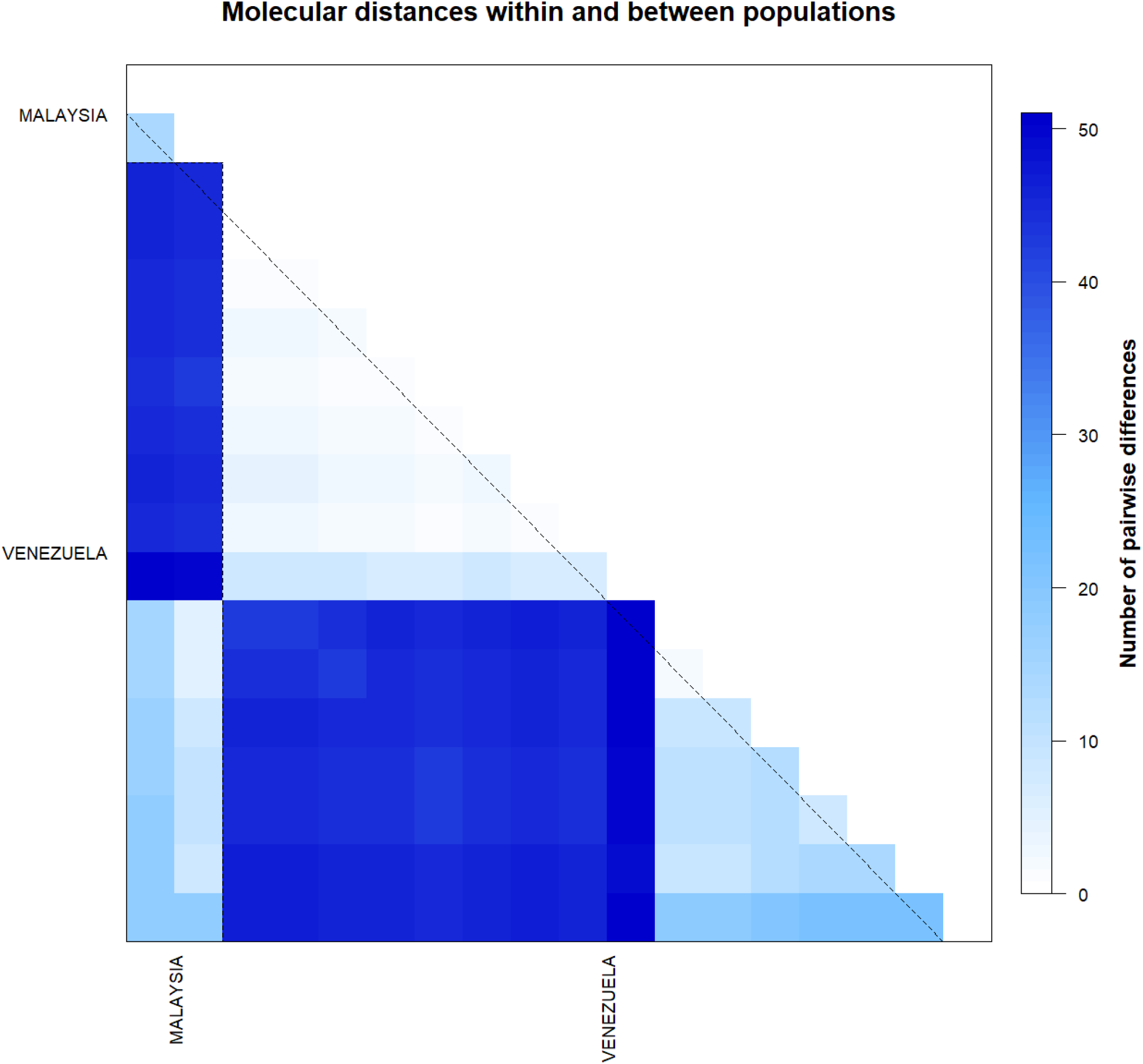
Matrix of inter haplotypic distance and number of polymorphic sites in the sequences of SARS-CoV-2 carriers of the D614G mutation. **Note the great variation between haplotypes from Venezuela**. *Generated by the statistical package in R language using the output data of the Software Arlequin version 3.5.1.2.

## Discussion

As the use of genetic diversity study methodologies not yet used in this customized PopSet for mutant SARS-CoV-2 genomes (D614G), it was possible to detect the existence of two distinct groups for the parcimonious-informative sites of viral genomes of Malaysia and Venezuela, containing significant inter-haplotypic variations only in the Venezuelan population. Perhaps, when other sequences from Malaysia are available in databases, this setting may change. The groups described here presented minimum patterns of structuring being slightly higher for the population of Venezuela, equaling the results obtained by Felix *et al*, 2020 for wild populations of SARS-CoV-2 in this same country. These data suggest that the relative degree of structuring present in Venezuela may be related to genetic flow. These levels of structuring were also supported by simple methodologies of phylogenetic pairing, such as UPGMA, which in this case, with a discontinuous pattern of genetic divergence between the groups (supports the idea of possible isolations resulting from past fragmentation events), especially when we observe a not so numerous amount of branches in the tree generated and with very few mutational steps.

These few mutations have possibly not yet been corrected by drift or lack of the founding effect, which accompanies the behavior of dispersion and/or loss of intermediate haplotypes over generations. The values found for the genetic distance support the presence of this continuous pattern of low divergence between the groups studied, since they considered important the minimum differences between the groups, when the haplotypes between them were exchanged, as well as the inference of values greater than or equal to those observed in the proportion of these permutations, including the p-value of the test.

The discrimination of the 18 genetic entities in the two localities was also perceived by large inter-haplotypic variations, hierarchical in all components of covariance: by their intra- and inter-individual differences or by their intra- and intergroup differences, generating a dendrogram that supports the idea that the significant differences found in the two countries, for example, were shared more in their form than in their number, since the result of estimates of the average evolutionary divergence found within these and other countries, even if they exist, are very low.

Based on the high level of haplotypic sharing perceived in Venezuela, tests that measure the relationship between genetic distance and geographic distance, such as the Mantel test, were dispensed with. The estimator φ, although extremely sensitive to any form of molecular variation (FU, 1997), supported the uniformity between the results found by all the methodologies employed, and can be interpreted as a phylogenetic confirmation that there is a consensus in the conservation of the mutant SARS-CoV-2 genome (D614G), in the two countries analyzed, and it is therefore safe to affirm that the small number of existing polymorphisms should not reflect major changes in the protein products of viral populations in the two countries. This consideration provides the security that, although there are differences in the haplotypes studied, these differences are minimal for both regions geographically analyzed and therefore it seems safe to extrapolate the levels of polymorphism and molecular diversity found in the samples of this study for other mutant genomes of SARS-CoV-2 in other countries, reducing speculation about a large difference between wild SARS-CoV-2 and mutant SARS-CoV-2 (D614G), at the level of protein products. Although the mutant form has a higher speed of transmission and infection, the analyses made in this study suggest small variations in protein products, especially in those that were targets of this study, bringing tranquility regarding the risk of death by COVID-19, as well as possible problems of adjustments to some molecular targets for vaccines.

